# Fecundity selection theory: concepts and evidence

**DOI:** 10.1101/015586

**Authors:** Daniel Pincheira-Donoso, John Hunt

## Abstract

Fitness results from the optimal balance between survival, mating success and fecundity. The interactions between these three components of fitness vary importantly depending on the selective context, from positive covariation between them, to antagonistic pleiotropic relationships when fitness increases in one reduce fitness of others. Therefore, elucidating the routes through which selection shapes life history and phenotypic adaptations via these fitness components is of primary significance to understand ecological and evolutionary dynamics. However, while the fitness components mediated by natural (survival) and sexual (mating success) selection have extensively been debated from most possible perspectives, fecundity selection remains considerably less studied. Here, we review the theory, evidence and implications of fecundity selection as a driver of sex-specific adaptive evolution. Based on accumulating literature on the life-history, phenotypic and ecological aspects of fecundity, we (*i*) suggest that ‘fecundity’ is restricted to refer to brood size per reproductive episode, while ‘annual’ and ‘lifetime fecundity’ should not be used interchangeably with ‘fecundity’ as they represent different life history parameters; (*ii*) provide a generalized redefinition of fecundity selection that encompasses any traits that influence fecundity in any direction (from high to low) and in either sex; (*iii*) review the (macro)ecological basis of fecundity selection (e.g., ecological pressures that influence predictable spatial variation in fecundity); (*iv*) suggest that most ecological theories of fecundity selection should be tested in organisms other than birds; (*v*) argue that the longstanding fecundity selection hypothesis of female-biased sexual size dimorphism (SSD) has gained inconsistent support, that strong fecundity selection does not necessarily drive female-biased SSD, and that this form of SSD can be driven by other selective pressures; and (*vi*) discuss cases in which fecundity selection operates on males.

## I. INTRODUCTION

Selection theory posits that fitness is a function of the balanced optimization between survival, mating success and fecundity (Williams, 1966; Rice, 2004; Charlesworth & Charlesworth, 2010). Three major mechanisms are responsible for the fitness dynamics that determine the trajectory of evolutionary change through differences in lifetime reproductive success (Darwin, 1859, 1871; Bell, 2008). Among these, natural and sexual selection explain how inter-individual differences in the exploitation of ecological resources and access to mates, respectively, regulates the genetic basis of phenotypic adaptations (Andersson, 1994; Schluter, 2000; Charlesworth & Charlesworth, 2010). Both mechanisms of selection form the central structure of current evolutionary theory, and have extensively been studied from a variety of conceptual and empirical perspectives (Williams, 1992; Andersson, 1994; Schluter, 2000; Gavrilets, 2004; Rice, 2004). The third mechanism, fecundity selection, traditionally describes the fitness advantages resulting from the selection for traits that increase fecundity (i.e., number of offspring; see section II) per reproductive episode (Roff, 2002). However, in contrast to the other two mechanisms of selection, fecundity selection has been the subject of considerably less research despite its major role in life-history theory (Lack, 1954; Williams, 1966; Shine, 1988; Roff, 2002), and its influence on the fitness outcomes of natural and sexual selection due to their interactions with the numbers of offspring that females produce (Sinervo, 2000; Ghalambor & Martin, 2001; Roff, 2002).

The hypothesis of fecundity selection was originally formulated by Darwin (1874) to explain the evolution of large body size in females, and in particular, the widespread evolution of female-biased sexual size dimorphisms (SSD) in which females are larger than males (Shine, 1988; Cox et al., 2003). The mechanistic basis of fecundity selection is that larger female size provides a greater body space to accommodate more offspring (Williams, 1966), and additionally, a higher capacity for energy storing to be subsequently invested into reproduction (Calder, 1984). The classical prediction of this hypothesis is that higher fecundity is a function of selection for larger female size, when fecundity depends on variation in female size and covaries positively with fitness. From this fundamental prediction a second major prediction can be derived: that strong fecundity selection generates directional selection on female body size or its components (see Braña, 1996; Scharf & Meiri, 2013), hence creating an asymmetrical selection effect between the sexes that drives female-biased SSD. Ever since Darwin (1874), most studies of fecundity selection have focused on overall female body size. However, a number of recent studies have expanded this view of fecundity selection towards a mechanism that influences fitness differentials via selection on any traits that are functionally linked to increased fecundity (e.g., Olsson et al., 2002; Parker et al., 2011; Winkler et al., 2012). In addition, it has been suggested that although fecundity selection can explain female-biased SSD, the expression and magnitude of this intersexual size asymmetry does not necessarily reflect the strength of fecundity selection (Zamudio, 1998; Olsson et al., 2002; Cox et al., 2003; Pincheira-Donoso & Tregenza, 2011; Soulsbury et al., 2014). Therefore, female-biased SSD can evolve in the absence of fecundity selection (e.g., via sexual selection for smaller male body size), and this form of selection can, in turn, be strong in the absence of female-biased SSD (e.g., when sex-specific directional selection pressures operate independently in both sexes). Finally, the rather dogmatic view that fecundity selection is mostly, or entirely, a female-specific mechanism has been relaxed by studies suggesting that variation in male body size or body shape components can have a causal effect on fecundity (Savalli & Fox, 1998; Hoffman et al., 2006; Wilson, 2009; Winkler et al., 2012).

Collectively, the fundamental assumptions and tenets that have characterized fecundity selection since its formulation demand the need for important modifications to this original concept, given the emergence of new perspectives and evidence that reveal a much more complex evolutionary force in terms of its targets (i.e., fecundity-fitness relationships, traits, sexes). However, despite the remarkable role of variation in fecundity among organisms of the same and different species, and the underlying selection behind the traits that regulate this variation, no review on the subject has appeared. In this paper, we review the concept of fecundity selection and its implications in evolutionary biology and ecology. We discuss the usage of central terms (such as ‘fecundity’) framed within life history theory, the impact of fecundity selection on traits other than body size and on males, and the (macro)ecological basis and consequences of this form of selection as the basis of major events of evolutionary change.

## II. THE CONCEPTUAL FOUNDATION OF FECUNDITY SELECTION

Fecundity selection has historically been regarded as the mechanism that promotes evolution of larger brood sizes via evolution of larger female body size (e.g., Cox et al., 2003; Fairbairn et al., 2007; Pincheira-Donoso & Tregenza, 2011). In its original formulation as the ‘fecundity advantage hypothesis’, Darwin (1874: 332) stated that ‘*increased size must be in some manner of more importance to the females…and this perhaps is to allow the production of a vast number of ova*’. Darwin (1874) derived the fecundity selection hypothesis to explain, in part, the evolution of female-biased sexual size dimorphism across animal species (section IV). The idea that fecundity selection targets female body size has implicitly prevailed in the literature given the overwhelming volume of empirical evidence revealing a positive relationship between female size and fecundity (Shine, 1988; Reiss, 1989; Stearns, 1992; Roff, 2002; Blanckenhorn, 2005; Fairbairn et al., 2007), especially amongst ectotherms, e.g., insects (Honek, 1993; Preziosi et al., 1996), fish (Wootton, 1979; Morita & Takashima, 1998; Foster & Vincent, 2004; Wilson, 2009), and reptiles (Cox et al., 2003; Cox et al., 2007; Stephens & Wiens, 2009; Pincheira-Donoso & Tregenza, 2011; Meiri et al., 2012). This relationship, however, is less robust in birds and mammals (Boyce, 1988; Shine, 1988; Purvis & Harvey, 1995; Lindenfors et al., 2007; Szekely et al., 2007). Importantly, this positive relationship between body size and fecundity is due to the fact that overall body size may often correlate positively with body regions that directly influence variation in fecundity, and not necessarily because body size is as a whole the responsible for levels of fecundity. For example, a positive body size-fecundity relationship can result from a positive relationship between female abdomen volume (where embryos are actually allocated) and body size, when fecundity is a direct function of available abdominal space (Andersson, 1994; Braña, 1996; Olsson et al., 2002; Scharf & Meiri, 2013). Therefore, a first needed step is to abandon the definition of fecundity selection as an overall body size-specific form of selection, and instead conceive it as a force that selects for any phenotypic trait(s) that increases fitness through fecundity.

Although far less abundant than examples involving overall female size, cases identifying fecundity selection on specific body regions that increase fecundity have started to emerge. For example, Olsson et al. (2002) showed that positive directional fecundity selection in a lizard (*Niveoscincus microlepidotus*) targets female trunk length for increased fecundity, while negative directional sexual selection targets male trunk length. Despite the strong positive effect of fecundity selection on female trunk length, males are significantly larger in body size (i.e., male-biased SSD). Therefore, this study supports the fecundity, but not the body size (or SSD), prediction of fecundity selection. Obviously, in contrast to animal-centred research, this is not a problem for fecundity selection studies on plants, where fecundity differentials are linked to a number of structural reproductive traits other than overall plant size (e.g., Stewart & Schoen, 1987; Shaanker et al., 1988).

A recurrent issue in fecundity-related studies is the usage of the term fecundity to refer to different forms of reproductive output. In the current literature, ‘fecundity’ is interchangeably employed to refer to the number of offspring per brood, per breeding season, and/or during a female’s lifetime. However, these measures of reproductive output represent importantly different life history parameters that can influence fitness in fundamentally different ways, given that fecundity and fitness are not always equivalent (e.g. Williams, 1966; Shine, 1988; Roff, 2002; see section II.1). In fact, the foundation of Williams’s (1966) central life history principle that describes trade-offs between current and future reproduction relies on the explicit distinction between current and future reproductive output (section II.1). Similarly, offspring per breeding season is also a parameter with a different meaning given that some species with high fecundity per brood can have low reproductive output per breeding season (e.g., one large clutch a year), while other organisms, such as *Anolis* lizards, lay a one-egg clutch multiple times per season (Losos, 2009). Therefore, we suggest the use of ‘lifetime fecundity’ (LF) for fecundity during an individual’s lifetime (and which will sometimes be the same as lifetime reproductive success), and ‘annual fecundity’ (Badyaev & Ghalambor, 2001) for fecundity per breeding season. ‘Fecundity’ should be *restricted* to refer to the number of offspring produced per brood per single reproductive episode (e.g., eggs in one single clutch). This distinction may prove operationally useful, and we recommend that this terminology is adopted in future studies. Therefore, strictly speaking, none of these definitions of fecundity should (from a conceptual point of view) be treated as equivalent to fitness, which we define as number of offspring that survive to breed (Lack, 1954; Fairbairn, 2006; Hunt & Hodgson, 2010). However, from a practical point of view, we refer to fitness as LRS (Fairbairn, 2006).

### (1) Fecundity and lifetime reproductive success

Differential fitness within a population can often be a function of differential fecundity. The most straightforward link between fitness and fecundity is the positive relationship between the two, such that phenotypes that confer higher fecundity are favoured by selection due to their fitness advantage (Rockwell et al., 1987; Stewart & Schoen, 1987; Godfray et al., 1991; Roff, 2002). When fecundity correlates positively with fitness, selection takes the shape of a positive directional fitness function. Positive directional selection is often a central idea implicit in studies of fecundity selection, which has also been repeatedly satisfied by empirical observation (e.g., Rockwell et al., 1987; Gibbs, 1988; Godfray et al., 1991). However, life history theory predicts that selection favours traits whose expression enhances lifetime reproductive success (LRS). And, given that fitness is not necessarily a constant positive function of fecundity, higher fecundity does not necessarily reflect higher fitness (Williams, 1966; Shine, 1988; Godfray et al., 1991; Roff, 2002).

Fecundity can compromise fitness via two main antagonistic interactions described by two central life history principles, namely, Lack’s (Lack, 1947) and Williams’s (Williams, 1966) principles. Both principles rely on optimality models which predict that fecundity is described by a negative quadratic (stabilizing) selection fitness function where intermediate fecundity values result in higher reproductive success (Williams, 1966; Boyce & Perrins, 1987; Gustafsson & Sutherland, 1988; Stearns, 1992; Sinervo, 2000). First, Lack’s principle predicts that selection favours brood sizes that yield the highest number of viable offspring that survive to breed (Lack, 1947, 1954). Hence, it describes a trade-off between offspring quantity and quality (Smith et al., 1989; Sinervo et al., 1992; Sinervo, 2000; Roff, 2002; Sibly et al., 2012). Second, Williams’s principle incorporates the costs of parental energy allocation into reproduction to predict that the extent of investment in current reproduction entails a reduction in future reproduction, and hence, compromises LRS (Williams, 1966; Nur, 1984; Gustafsson & Sutherland, 1988; Sinervo & DeNardo, 1996; Roff, 2002). Consequently, both principles predict that the relationship between fitness components (offspring and parental viability and/or future fecundity) and fecundity is consistently under a strong trade-off with natural selection arising from the costs of reproduction (Gustafsson & Sutherland, 1988; Sinervo, 2000; Ghalambor & Martin, 2001; Roff, 2002). Therefore, although fecundity selection by historical definition implies selection favouring higher fecundity, this form of selection is inevitably countered by natural selection on lifetime reproductive success through viability, thus often creating non-linear fitness functions.

### (2) The shape and definition of fecundity selection

A singular aspect of fecundity selection is that, strictly speaking, fitness functions cannot be obtained by regressing a measure of fitness on a phenotypic trait, given that fecundity (as defined in the previous subsection) is a measure of phenotype rather than a measure of fitness. Thus, fecundity is the trait on which fitness (e.g., LRS) should be regressed. As explained above, the reason for this is that fitness is a function, rather than an equivalent, of fecundity (e.g., see Williams’s principle). A fundamental implication that emerges from this fitness-fecundity relationship is that the original formulation of fecundity selection (Darwin, 1874) fails to capture the multiple effects that fecundity exert on fitness by defining it as a selective force with positive directional effects. For example, the Williams’s principle predicts that high fecundity can have a fitness disadvantage (Williams, 1966; Shine, 1988; Roff, 2002). Also, other studies show that low fecundity (although not necessarily low annual fecundity) can be selected for when small broods have a fitness advantage. For example, the bet-hedging fecundity strategy (i.e., adaptive reduction of brood size) has been demonstrated across a wide diversity of organisms (Gillespie, 1974; Lehmann & Balloux, 2007; Griebeler et al., 2010). This phenomenon evolves, among other causes, when predation intensity is high and mothers spread the risk of offspring loss across multiple independently laid smaller clutches (Griebeler et al., 2010; see also section III.1.b). As indicated above, the lizards of the prolific *Anolis* radiation, which lay multiple one-egg clutches yearly (Losos, 2009), are a prime example of this phenomenon. Therefore, selection for fecundity is known to favour different brood sizes depending on the way that environmental demands influence fitness, and hence, the traditional definition of fecundity selection describes only the positive extreme of the whole range of outcomes that result from this mechanism.

Consequently, the above discussion demands a redefinition of the concept of fecundity selection. Here, we define fecundity selection as ‘*differential lifetime reproductive success (i.e., fitness) as a function of variation in phenotypic traits that influence fecundity*’. Our definition (*i*) generalizes the action of fecundity on reproductive success by incorporating the fitness advantages that can result from selection for reduced fecundity; (*ii*) does not restrict the action of fecundity selection to female body size, but instead, recognizes that any heritable phenotypic trait that results in enhanced fitness through adjustments in fecundity can adapt via fecundity selection; (*iii*) removes from the concept the predominant view that fecundity selection operates on females only (as will be discussed in section VI, male phenotypes influence fecundity in species where males play a substantial role in embryo development). Therefore, collectively, our definition denies presumptions on the trait- and sex-specificity of this mechanism. Another advantage of our definition is that it does not restrict the action of fecundity selection to animals only. Fecundity selection in fact operates on plants (e.g. Clegg & Allard, 1973; Lloyd, 1987; Stewart & Schoen, 1987; Shaanker et al., 1988; Vekemans et al., 1998; Cummings et al., 2002).

## III. THE (MACRO)ECOLOGY OF FECUNDITY SELECTION

The contribution of fecundity to fitness is influenced by multiple ecological factors that impose limits to the evolution of brood size via the costs that production of offspring per reproductive episode exerts on reproductive success. A major implication of this interaction between fecundity and ecological pressures is that fecundity selection is not necessarily positive and directional, but can maximize fitness via adaptive reductions of brood size depending on the ecological context.

Ecological pressures on fecundity arise from a number of environmental components, including climate (e.g., seasonality, temperature, and a number of factors that vary with geographic gradients), and from interspecific interactions, such as predation (Williams, 1966; Godfray et al., 1991; Roff, 2002). For example, extensive evidence shows that predation has a major impact on fecundity (e.g., larger broods can be more vulnerable to predators when multiple offspring are more visited by caring parents or produce more noise). Interestingly, given that most (if not all) of these ecological pressures vary with geography, the analysis of such factors in a macroecological perspective can shed light on the general factors underlying patterns of variation in fecundity.

### (1) Fecundity in a macroecological context: the ‘Moreau-Lack’s rule’

Spatial variation in selection arising from environmental differences along geographic gradients exerts major effects on local life history adaptations, which results in the expression of large-scale patterns of life history evolution (Moreau, 1944; Stearns, 1992). One of the most important such macroecological life history generalizations is the observation that fecundity increases predictably with increasing latitude (Moreau, 1944; Lack, 1954; Bennett & Owens, 2002). This hypothesis, proposed by Moreau (1944) and Lack (1954) to explain bird life histories, has achieved the status of a life history paradigm (e.g., Musvuugwa & Hockey, 2011), for which we propose the name ‘Moreau-Lack’s rule’. Historically, this rule has heavily focused on clutch size variation in birds (e.g., Jetz et al., 2008; McNamara et al., 2008a; Griebeler et al., 2010).

During the early foundation of Moreau-Lack’s rule, Moreau (1944) suggested that the observed differentials in fecundity along geographic gradients were unlikely to be explained by one single factor. However, he suggested that higher mortality in more seasonal environments (i.e., higher latitudes) might be a predominant factor behind this life history pattern. Later, Lack (1954) conferred primary importance (for both offspring and parent fitness) to the effect of food availability and management across different latitudes. He suggested that day-length increases toward the poles (e.g., in ∼50% between the equator and central Europe, and nearly 100% between the former and the arctic) result in increased opportunities for parents to collect more food. Thus, the conditions to successfully produce more offspring are better (Lack, 1954). However, Lack claimed that extreme day-lengths (in the poles) may be detrimental for parental survival as permanent food searching would exhaust them. Additionally, he suggested that declines in food availability in extremely high latitudes (e.g., Scandinavia) may drive the expression of a similar fecundity-latitude function closer to the poles. Therefore, this would create a non-linear gradient with increases toward moderately high latitudes, but then a decrease from these latitudes toward the poles (Lack, 1954). For decades, the prediction of a macroecological fecundity gradient has consistently been demonstrated by numerous bird studies conducted both within (Klomp, 1970; Koenig, 1984; Young, 1994; Koenig & Gwinner, 1995; Sanz, 1998; Dunn et al., 2000) and among species (Martin, 1996; Bohning-Gaese et al., 2000; Martin et al., 2000; Cardillo, 2002; Martin, 2002; Jetz et al., 2008; Rubolini & Fasola, 2008). Likewise, similar patterns have been shown to hold in a number of other organisms, including mammals (Lord, 1960; Jackson, 1965; Conaway et al., 1974; Innes, 1978; Cockburn et al., 1983; Swihart, 1984; Whorley & Kenagy, 2007; Bywater et al., 2010), reptiles (Fitch, 1970; Iverson et al., 1993; Forsman & Shine, 1995; Litzgus & Mousseau, 2006; Simoncini et al., 2009), amphibians (Kuramoto, 1978; Tilley, 1980; Cummins, 1986; Morrison & Hero, 2003), and fish (Fleming & Gross, 1990; Kokita, 2004). The consistency of this latitudinal fecundity gradient (LFG, hereafter) has, as indicated above, achieved the status of a life history paradigm (Musvuugwa & Hockey, 2011; Rose & Lyon, 2013). However, as can be expected for any macroecological generalization, other studies have also failed to support this rule in a number of cases, including vertebrates and invertebrates (e.g., Yom-Tov et al., 1994; Jansen et al., 2014).

The accumulation of empirical evidence and the development of more complex and realistic theoretical models (McNamara et al., 2008a; Griebeler et al., 2010) have resulted in important progress for understanding the shape of, and the proximate mechanisms underlying, this fecundity gradient. First, apart from the major conclusion that Moreau-Lack’s rule is a consistent macroecological pattern, another conclusion that can be drawn from these studies is that the non-linear gradient expected by Lack is not a generality. Indeed, the studies based on the largest numbers of species from the highest diversities of areas in the world (Cardillo, 2002; Jetz et al., 2008) have shown a consistent increase of clutch size with elevation. Therefore, these observations (still heavily bird-biased) seem to rule out Lack’s (1954) non-linear latitudinal expectation. Second, this evidence has led to the consolidation of four main hypotheses which, probably more dependently than independently, can explain Moreau-Lack’s rule. We present these hypotheses below.

#### (a) Seasonality and Ashmole’s hypothesis

Increasing latitudes result in increasing seasonality, which determines patterns of temporal fluctuation of resources throughout the year. Based on this premise, Ashmole (1963) suggested that (bird) fecundity varies spatially as a function of spatial patterns of seasonality. Toward higher latitudes, where differences in food availability between seasons are greater, birds are predicted to suffer higher mortality rates during the winter as a result of lower resource abundance. These declines in population density turn into favourable feeding conditions for survivors, as the per capita food availability is higher when the next breeding season starts, which therefore facilitates the opportunities for more energy to be invested in fitness through higher fecundity. Ricklefs (1980) later made the distinction that latitudinal fecundity gradients may result specifically from factors that reduce population density during the winter (the non-reproductive season) rather than by higher relative abundance of food during the start of the breeding season. Thus, Ashmole’s hypothesis relies on resource availability as a driving factor behind density-dependent dynamics of fecundity. Decades of both theoretical (McNamara et al., 2008a; Griebeler et al., 2010) and empirical confirmation (Ricklefs, 1980; Koenig, 1984; Birt et al., 1987; Dunn et al., 2000; Yom-Tov & Geffen, 2002; Jetz et al., 2008) have made this hypothesis potentially the main mechanistic explanation for the pattern described by Moreau-Lack’s rule.

However, an alternative hypothesis for the latitudinal fecundity gradient under the effect of seasonality exists: longer days during the breeding season at higher latitudes give parents the opportunity to spend more time foraging each day, which thus increases daily total food delivery to the brood, and hence, increases the potential to sustain more offspring (Rose & Lyon, 2013). Yet, despite the mechanistic appeal of this hypothesis – which was questioned by Lack (1954) himself – it remains virtually untested. Interestingly, a recent study conducted on the latitudinally widespread tree swallow (*Tachycineta bicolor*), revealed support for this ‘day length’ hypothesis, while showing little support for the classical alternative (Rose & Lyon, 2013). Thus, these authors suggested that the length of an animal’s workday can be an important, yet unappreciated, factor behind the evolution of fecundity in a macroecological context. However, once again, a major limitation for the generality of these hypotheses is the lack of studies in organisms other than birds. So far, only a few studies conducted in non-avian organisms have attempted to address aspects of the Ashmole’s hypothesis (e.g., Lindstedt & Boyce, 1985; Sainte-Marie, 1991; Iverson et al., 1993), although not always focusing on its links to fecundity.

#### (b) Differences in nest predation

Nest predation exerts major effects on offspring and adult fitness, and hence, spatial gradients of nest predation are expected to cause fecundity gradients (Martin, 1996; McNamara et al., 2008a; Lima, 2009; Griebeler et al., 2010). Traditionally, it has been assumed that larger clutches/litters are more detectable by predators as a result of more frequent parental care (e.g., feeding visits) and overall noisier offspring (Skutch, 1949; Slagsvold, 1984; Godfray et al., 1991; Griebeler et al., 2010). Therefore, areas with intense nest predation are predicted to select for multiple nests of smaller clutches with shorter development times (Skutch, 1949; Martin, 1996). Given that predation is thought to intensify toward the tropics (e.g., Griebeler et al., 2010), the consistent observation that females tend to produce multiple smaller clutches in these environments has been explained as a consequences of intense nest predation. However, although nest predation is often invoked as a potentially major driver of the latitudinal fecundity gradient (Skutch, 1949; Slagsvold, 1982; Lima, 1987; Kulesza, 1990; Jetz et al., 2008; Martin & Briskie, 2009), this hypothesis has also often been discredited. For example, intense nest predation can slowdown developmental rates because reduced parental visits result in food limitation (Ghalambor & Martin, 2001), and development, at least in birds, is often slower toward the tropics (Martin, 1996; Geffen & Yom-Tov, 2000). Also, this hypothesis implicitly assumes that predation is caused by visually-oriented predators, despite the fact that chemically-oriented predators (those with highly developed olfactory senses, such as snakes) are highly common (Thompson & Burhans, 2003). Not surprisingly then, evidence for the plausibility of this hypothesis is conflicting. Both mathematical models (McNamara et al., 2008a; Griebeler et al., 2010) and empirical analyses (Geffen & Yom-Tov, 2000; Robinson et al., 2000; Martin et al., 2006; Biancucci & Martin, 2010) have often suggested that nest predation appears to play only a secondary (if any) role. Others reveal more complex interactions between clutch size and predation. For example, Biancucci & Martin (2010) showed that nest predation risk increases with nest size. However, nest size is not necessarily related to clutch size and hence, nest predation risk does not, according to this view, drive the latitudinal fecundity gradient. Finally, differentials in nest predation intensity have been found to be unimportant in explaining elevational fecundity gradients (e.g. Badyaev, 1997; Badyaev & Ghalambor, 2001), or have resulted in conflicting observations (Lu, 2008). Given the overwhelming tendency for research on birds, studies on non-avian models are needed to assess the generality of this hypothesis.

#### (c) *Length of breeding season hypothesis* (LBS).

Populations from higher latitudes experience reduced opportunities for multiple reproductive episodes per year as a result of increasing seasonality and shorter warm seasons. Therefore, according to the LBS hypothesis, the intensity of fecundity selection increases as a function of increasing latitudes (Lack, 1954; Fitch, 1970; Tinkle et al., 1970; Cox et al., 2003; Pincheira-Donoso & Tregenza, 2011). The primary prediction of this hypothesis is that shorter breeding seasons create fecundity selection for larger clutches that compensate for reduced reproductive frequency. The specific factors linking length of the warm season with reduced reproductive frequency can be common to all organisms or vary among lineages. For example, factors such as prolonged snow cover or delayed emergence of food are expected to delay the start of the breeding season, thus reducing reproductive opportunities for organisms in general (Cox et al., 2003; Griebeler et al., 2010). In contrast, thermal constraints on annual and daily activity can cause more severe effects on ectotherms. In high latitudes (and elevations), not only environmental temperatures are lower, but also warm hours per day are fewer (Nagy & Grabherr, 2009), which makes reproductive seasons even more severely limited for ectothermic organisms. Therefore, compared to endotherms, ectotherms have even more limited opportunities for reproduction (Fitch, 1970; Tinkle et al., 1970; Fitch, 1978, 1981; Cox et al., 2003; Cox et al., 2007; Pincheira-Donoso & Tregenza, 2011; Pincheira-Donoso et al., 2013). As a result, theory predicts that fecundity selection favours traits that maximize brood size (e.g., female body size) in each of these infrequent reproductive episodes, so that reduced opportunities for reproduction are compensated by higher fecundity (Godfray et al., 1991; Cox et al., 2003; Pincheira-Donoso & Tregenza, 2011). On the other hand, long breeding seasons (toward the tropics) are expected to select for smaller clutches as females spread the energetic costs of brood production over time and the risk of mortality by predation across space (Griebeler et al., 2010; section III.1.d). In a modelling-based study, Griebeler et al. (2010) investigated the factors that explain variations in avian fecundity along latitudinal gradients, focusing on the effects of resource seasonality, nest predation, and LBS. Their models revealed that LBS in combination with resource seasonality and predation explain the observed trends in clutch size, i.e., higher fecundity toward the poles. However, resource seasonality is the predominant factor (it can drive the latitudinal trend by itself), while LBS was found to be a weak independent predictor of gradients in fecundity and annual fecundity.

Phylogenetic tests of the LBS hypothesis have primarily been performed on reptile models. In squamates (lizards and snakes) in particular, the breeding season effect is expected to be intensified by viviparity, which compromises reproductive frequency through prolonged embryo retention within the mother until development is completed (Blackburn, 2000; Cox et al., 2003; Shine, 2005). This constraining effect is further reinforced as viviparity predominantly evolves among species occurring in cold climates (Lee & Shine, 1998; Hodges, 2004; Shine, 2005; Schulte & Moreno-Roark, 2010; Pincheira-Donoso et al., 2013), where oviparity is extremely rare (Shine, 2005; Pincheira-Donoso et al., 2013). A traditional approach for linking the intensity of fecundity selection with shorter breeding seasons consists in investigating variation in the degree of female-biased SSD as a function of, for example, increasing latitude (or elevation). Therefore, selection is predicted to favour early reproduction and higher brood number in species that reproduce frequently, while delayed reproduction accompanied by higher fecundity is predicted in those that reproduce infrequently (Tinkle et al., 1970; Fitch, 1978, 1981; Cox et al., 2003).

Two phylogenetic studies have tested this prediction. In a global-scale analysis (covering 302 species from 18 families), Cox et al. (2003) investigated the relationship between multiple proxies for reproductive frequency and SSD as a classical indicator of fecundity selection intensity (see section IV, for discussion). This study supported the central predictions that female-biased SSD is significantly correlated with clutch size, reproductive frequency (i.e., number of broods per season), and reproductive mode (i.e., oviparity/viviparity). However, tests using the length of reproductive season (i.e., from earliest observed vitellogenesis to latest oviposition) and latitude as independent proxies for the strength of fecundity selection revealed no relationship with SSD. More recently, Pincheira-Donoso & Tregenza (2011) investigated the effect that an extreme geographic gradient (combining latitudinal/elevational climatic effects) exert on variation in fecundity among species of the prolific *Liolaemus* lizard radiation. Consistent with Cox et al.’s (2003) study, these authors failed to observe the predicted negative relationship between LBS and fecundity. However, Pincheira-Donoso & Tregenza (2011) argued that these findings may in fact indicate that the strength of fecundity selection increases as LBS decreases. Given that colder climates entail further costs to fecundity, such as shorter and unstable days (Pincheira-Donoso et al., 2008; Nagy & Grabherr, 2009), and longer gestation via viviparity (Shine, 2005; Meiri et al., 2013; Pincheira-Donoso et al., 2013), smaller brood sizes would in fact be expected in these short breeding season environments. Thus, similar brood sizes between warm and cold environment species may actually imply that stronger fecundity selection compensates this otherwise expected decline in fecundity under such adverse conditions. Therefore, collectively, evidence supporting the LBS hypothesis remains elusive. Empirical tests are, however, still very few and the taxonomic coverage of model organisms greatly limited too, which leaves a clear open niche for future research.

#### (d) The ‘bet-hedging strategy’ hypothesis

There is a tendency for the selective demands underlying macroecological fecundity gradients to be decomposed into the three above factors (i.e., food seasonality, nest predation and LBS). According to Griebeler et al. (2010), this ‘divisive’ approach may have limitations as it assumes that these factors operate independently, while the three can in fact operate simultaneously to drive the gradient. Thus, Griebeler et al. (2010) proposed a new integrative hypothesis to explain the latitudinal fecundity gradient that combines the above three sources of natural selection. This hypothesis relies on the observation that food availability determines the total annual fecundity, which in turn depends on the LBS (and which in turn determines numbers of breeding attempts per season), and that nest predation determines how this total annual fecundity is spread over different broods (e.g., Gillespie, 1974; Zanette et al., 2006; Lehmann & Balloux, 2007; Lima, 2009; Griebeler et al., 2010). Griebeler et al. (2010) suggested that (*i*) the ‘empty slots’ caused in populations by winter mortality (i.e., Ashmole’s hypothesis), determine the success of broods in relation to brood size (i.e., fecundity), and that (*ii*) brood size, in turn, is a function of nest predation risk and LBS. Therefore, where food seasonality is low, LBS is long and predation intensity is high (as in tropical latitudes), selection favours a ‘bet-hedging’ strategy such that the risk of predation is spread over multiple smaller clutches (Yasui, 1998; Farnsworth & Simons, 2001; Griebeler et al., 2010), and hence, the success of number of broods relative to their size increases. In colder climates, at higher latitudes, this strategy is neither viable given that LBS are short, nor selected for given that nest predation is low. Therefore, this hypothesis predicts fewer and larger broods toward higher latitudes as a result of the functional interaction among the above three factors (Griebeler et al., 2010). Despite this hypothesis offers an explicitly integrative view, it is not an untested set of predictions as, in fact, all its components have been investigated within the framework of the above three hypotheses. Hence, the bed-hedging hypothesis is mostly a summary hypothesis than a new hypothesis. Consequently, evidence for and against the specific components of this hypothesis comes from a number of studies that have been presented in this section which deal with the components of Griebeler et al.’s (2010) hypothesis (e.g., Ricklefs & Bloom, 1977; Slagsvold, 1982, 1984; Zanette et al., 2006; Jetz et al., 2008).

The bet-hedging prediction in a macroecological context is not, however, supported in at least a fish species. In the pipefish *Syngnathus leptorhynchus*, characterized by male-pregnancy, reproductive success is optimized by adjustments of life-history parameters along the latitudinal gradients it occupies in North America (Wilson, 2009). One such adjustment involves the strategy of female egg transference to males. In contrast to the predicted pattern, female bet-hedging increases toward higher latitudes, where eggs are distributed among more males (Wilson, 2009). It has been interpreted that this strategy evolves in response to harsher environments where females aim to reduce costs of mortality by spreading the probability of mortality across multiple males.

### (2) The fecundity gradient paradox

An important underlying generalization of the original formulation of Moreau-Lack’s rule, and common to most subsequent studies on it, is that the fecundity gradient is driven by environmental variation in latitude. Hence, it can be expected that the environmental mechanisms that operate on life history variation along latitudes (e.g., seasonality, length of breeding season, food availability, temperature) should operate in similar ways along elevational gradients, creating an altitudinal fecundity gradient (AFG). In fact, a number of studies have investigated fecundity differentials along altitudinal gradients, and as in other related fecundity phenomena, the focus is heavily biased toward birds. Remarkably, however, elevational fecundity variation strongly opposes the latitudinal pattern. With substantial consistency, fecundity decreases with increasing elevation (Badyaev, 1997; Badyaev & Ghalambor, 2001; Lu, 2005; Sandercock et al., 2005; Johnson et al., 2006; Jin & Liu, 2007; Kleindorfer, 2007; Lu et al., 2008; Lu et al., 2009; Lu et al., 2010; Lu, 2011; Ramirez-Bautista et al., 2011), and hence, colder and more seasonal environments appear to select for smaller brood sizes. Other studies, however, have revealed no fecundity differentials along altitudes (Lu, 2008; Bears et al., 2009), or larger clutches toward higher elevations (Carey et al., 1982; Martin et al., 2009; Camfield et al., 2010). Despite this remarkable asymmetry, similar environmental factors have traditionally been employed to explain the expression of each (latitudinal and altitudinal) fecundity gradient separately. Among them, continuous variation in seasonality of resources, nest predation intensity, length of breeding seasons, and decreases in environmental temperatures are regarded as the major sources of selection. Interestingly, these factors have in common their continuous variation in similar directions with increasing latitude and elevation. For example, risk of nest predation has commonly been noted to decrease towards higher latitudes and elevations, as well as environmental temperatures and the length of breeding seasons do. Therefore, the evolution of these two fecundity gradients in opposite directions under (apparently) similar gradients of selection based on the above factors suggests an appealing life history paradox.

The opposite responses of fecundity gradients to latitudinal and elevational gradients suggests that, (*i*) the selective effects of the above factors interact in different ways with other selective factors that differ between geographic gradients in latitude and in elevation (e.g., geographic clines in atmospheric conditions), to create different forms of net multivariate selection on fecundity strategies; and (*ii*) the evolutionary scenarios offered by geographic variation across latitudes provides opportunities and constraints to life history adaptations that differ from the scenarios offered by elevational variation, causing opposing life history responses to selection. For example, ecological and climatic zones are replaced in considerably shorter distances as elevation increases compared to latitude (Bonan, 2008). The most likely explanation may involve a combination of both factors.

A major potential driver of this fecundity paradox might be spatial gradients in atmospheric oxygen concentrations. It is broadly known that multiple climatic factors (e.g., temperature) vary in the same direction with increasing latitude and elevation (Bonan, 2008; Nagy & Grabherr, 2009). In contrast, given that oxygen availability declines as a function of decreasing atmospheric pressure (Nagy & Grabherr, 2009), its concentrations steeply decline with elevation, but do not decline with latitude (Nagy & Grabherr, 2009). In addition, spatial changes in oxygen concentrations are known to exert major impacts on egg development. For example, it has consistently been shown in reptiles (Deeming & Ferguson, 1991; Kam, 1993; Warburton et al., 1995; Andrews, 2002; Deeming, 2004) and birds (Black & Snyder, 1980; McCutcheon et al., 1982) that low levels of oxygen concentrations reduce developmental success. Aspects such as embryonic differentiation and growth rates, water uptake, duration of incubation, growth of the chorioallantonic membrane, egg survival, and hatchling size are known to be negatively affected by hypoxia (Andrews, 2002; Parker et al., 2004). Among *Sceloporus* lizards, for example, successful development depends on high levels of *in utero* oxygen (Andrews & Rose, 1994; Andrews, 2002; Parker et al., 2004; Parker & Andrews, 2006). Also, it has been suggested that transitions from oviparity to viviparity in reptiles may be promoted by declines in atmospheric oxygen (Hodges, 2004; Lambert & Wiens, 2013), and that in turn, viviparity reduces reproductive opportunities, thus promoting larger clutches (Cox et al., 2003; Pincheira-Donoso & Tregenza, 2011). Finally, it has been observed that development can be compromised in larger clutches given that oxygen concentrations decline towards their centre (Ackerman, 1977; Ackerman & Lott, 2004). Moreover, rates of oxygen declines versus CO_2_ increases in turn increase with clutch size (Ackerman & Lott, 2004). Consequently, all these factors taken together suggest that fecundity may vary differently along elevational and latitudinal gradients in response to the effects that differential gradients of oxygen have on the developmental success of eggs.

Alternatively, the paradox may be the result of incorrect generalizations about the geographic variation in the factors predicted to cause it. For example, as stated above (section III.1.b), smaller clutches are predicted in high-predation environments, and predation intensity tends to decline with latitude. However, predation does not necessarily decline with elevation. For instance, while clutch size can increase with elevation when predation declines towards higher altitudes (Camfield et al., 2010), higher predation towards higher altitudes has been found to promote smaller clutches with increasing elevation (Sandercock et al., 2005; Lu, 2011). Therefore, although the *pattern* is inconsistent along the same geographic gradients (because the factors behind the patterns do not vary consistently with elevation), the *mechanistic effects* of the factors on fecundity clines are strongly consistent (i.e., smaller broods toward higher predation environments).

Whatever the best general explanation for both fecundity gradients, it seems clear that accounting for their expression is far more complex than currently appreciated by studies focusing separately on either of them. Indeed, this has been the consistent tendency, given that both the LFG and AFG are largely (and implicitly) treated as unconnected phenomena. Therefore, the mechanism underpinning this paradox remains an open question.

## IV. FECUNDITY SELECTION AND THE EVOLUTION OF SEXUAL DIMORPHISM

The evolution of sexual size dimorphism (SSD) is a complex phenomenon driven by the asymmetrical effect of intersexual antagonistic selection (IAS) operating differentially on the phenotype of males and female (Andersson, 1994; Blanckenhorn, 2005; Fairbairn, 2007; Cox & Calsbeek, 2009), or by their asymmetric non-genetic responses to context-specific environmental demands that influence body size development (Cox et al., 2003; Cox & Calsbeek, 2010). The adaptive evolution of SSD has traditionally been explained via sex-specific (mostly male-specific) positive directional sexual selection (Andersson, 1994; Cox et al., 2003; Fairbairn et al., 2007), disruptive natural selection (Shine, 1989), directional fecundity selection on female traits (Darwin, 1874; Cox et al., 2003), or a combination of different mechanisms of selection operating in opposing directions between the sexes (Zamudio, 1998; Olsson et al., 2002; Pincheira-Donoso & Tregenza, 2011). The classical hypothesis of fecundity selection-driven SSD was originally formulated by Darwin (1874) to explain the evolution of female-biased SSD. According to this hypothesis, fecundity selection promotes increased fecundity through a directional effect on female body size in species where reproductive output is compromised (see section III for details). This female-specific effect is broadly assumed to create antagonistic selection on body size between the sexes, resulting in female displacements from males, and ultimately, in female-biased SSD (Cox et al., 2003; Pincheira-Donoso & Tregenza, 2011).

The fecundity selection hypothesis of SSD has extensively been invoked to explain the evolution of larger females in lineages where the direction (whether male- or female-biased) and magnitude of SSD vary among species. Indeed, the view that the magnitude of female-biased SSD reflects the strength of fecundity selection on female body size has become a traditional assumption of the fecundity selection hypothesis of SSD. Under this view, comparatively larger females indicate stronger intersexual antagonistic selection caused by stronger fecundity selection. For example, Shine’s (1988) main assumption when analysing the underlying rationale behind fecundity selection was, in fact, that females evolve larger body size than males when fecundity selection operates. However, the question is whether the expression and extent of female-biased SSD can be taken as a reliable signal of the action and strength of fecundity selection. A number of studies have suggested that the action of (even strong) fecundity selection on female body size does not necessarily translate into female-biased SSD (see below for details), or that this form of SSD may result from the antagonistic effects of selection mechanisms other than fecundity selection (Zamudio, 1998; Olsson et al., 2002; Pincheira-Donoso & Tregenza, 2011). In this latter case, theory suggests that larger females are not only the result of female-specific positive directional selection on body size, but can also be the result of negative directional selection on male size (Singer, 1982; Andersson, 1994; Zamudio, 1998). For example, Zamudio (1998) observed that female body size correlates positively with fecundity in *Phrynosoma* lizards with female-biased SSD. However, her phylogenetic evidence revealed that this form of SSD was more likely driven by negative directional selection on male size for earlier maturation. Similarly, Olsson et al. (2002) showed that strong female-specific fecundity selection does not alter pronounced male-biased SSD in a lizard (see next subsection for details). In another study (Wilson, 2009), a similar example was presented. Although body size in the pipefish *Syngnathus leptorhynchus* is sexually monomorphic, fecundity selection on male size (via male pregnancy; see section VI) has been suggested to be strong (Hoffman et al., 2006; Wilson, 2009). Interestingly, toward higher latitudes where waters become colder, male brood pouch volume increases via increases in male body size. This size gradient counterbalances the reduced opportunities for reproduction caused by extended brooding periods in cold climates and thus, maintains male reproductive success across the species range. Consequently, in line with other observations, fecundity selection does not drive SSD in these fish.

The problem with treating SSD as an indicator of fecundity selection is further revealed by perspectives in which fitness is broken up into its three fundamental components (survival, mating success and fecundity). Firstly, although (sexual) selection for mating success mostly predicts sex-specific effects on males, these effects can (as discussed above) select for smaller males, or in fact, for female traits when male mate choice operates (Andersson, 1994). Therefore, female-biased SSD can be driven by sexual selection. Second, the role for natural selection in the evolution of sexual dimorphism has increasingly been appreciated (Shine, 1989; Butler et al., 2000; Temeles et al., 2000; Losos et al., 2003; Fairbairn et al., 2007; Pincheira-Donoso et al., 2009; Temeles et al., 2010; Meiri et al., 2014). One central attribute of the ecological hypothesis of sexual dimorphism is that it does not predict a predominant direction in the evolution of SSD (i.e., whether it is mostly male-biased or female-biased). Instead, it only predicts that intersexual resource competition creates frequency-dependent natural selection for sexes to adapt to alternative ecological resources (e.g., Bolnick & Doebeli, 2003), and thus, sex-specific body sizes are pushed apart in the direction of any available niche space. Therefore, both males and females are in theory equally likely to evolve larger body size, or to remain monomorphic for size if other ecologically relevant (body shape) traits allow divergent access to different resources. Finally, although fecundity, natural and sexual selection can explain the evolution of female-biased SSD, a recent study showed that sexual selection is the predominant driver of sexual dimorphism across animals in general (Cox & Calsbeek, 2009). This finding, in line with previous evidence that sexual selection exceeds other selection mechanisms in the wild (Hoekstra et al., 2001; Kingsolver et al., 2001; Blanckenhorn, 2007), relegates the role of fecundity selection on sexual dimorphism to a secondary level. Remarkably, however, the role of natural selection was found to be even weaker, hence retaining fecundity selection as the second most important driver of SSD after sexual selection, which prevails as the primary driver (Cox & Calsbeek, 2009).

Collectively, although the role for fecundity selection in the evolution of female-biased SSD is broadly recognized (Cox et al., 2003; Shine, 2005; Cox & Calsbeek, 2009; Pincheira-Donoso & Tregenza, 2011), this form of SSD should not be treated as an indicator of the action or strength of fecundity selection. Thus, female-biased SSD does not necessarily imply the action of fecundity selection, while the absence of female-biased SSD does not imply that fecundity selection is not in operation (Singer, 1982; Zamudio, 1998; Olsson et al., 2002; Pincheira-Donoso & Tregenza, 2011). Similarly, in species in which fecundity selection operates on males (section VI), SSD is not a norm and hence, the same principle suggested above would expand into these situations of reversed sex-roles.

### (1) Sexually antagonistic fecundity selection on body regions

Sexual dimorphism is not the same as SSD. As argued earlier, the concept of fecundity selection has expanded from a body size-centred view to a more evolutionarily broad (and realistic) idea that involves any component of the phenotype that enhances fitness via adjustments in fecundity (section II). Fecundity selection is, therefore, predicted to target specific body regions rather than overall size, and when these effects are sexually antagonistic, sexual dimorphism for specific traits is likely to evolve (e.g., Olsson et al., 2002; Fairbairn et al., 2007; Winkler et al., 2012; Scharf & Meiri, 2013). A number of examples of sexual dimorphism presumed to be driven by fecundity selection exist. For example, in large-scale studies covering broad diversities of lizards, it has been found that fecundity selection targets female abdomen length specifically, where embryos are kept during development (Braña, 1996; Scharf & Meiri, 2013). Similarly, using quantitative genetic measures of sexually antagonistic selection, Olsson et al. (2002) revealed strong evidence for sex-specific directional fecundity selection on female abdomen length in the lizard *Niveoscincus microlepidotus*. Interestingly, male abdomen length was also found to be under selection, but negative directional sexual favouring shorter trunks, which are potentially advantageous for male-male contests (Olsson et al., 2002). Despite such clear effect of fecundity selection on females, male body size is considerably larger in this species (Olsson et al., 2002), which reinforces the view that fecundity selection can be strong, yet negligible in its role in driving female-biased SSD.

Additional examples of region-specific selection come from pipefish species (genus *Syngnathus*), in which sex roles are reversed and male-pregnancy occurs. In these fish, it has been shown that fecundity selection targets male- and female-specific body regions. Female pipefish transfer the eggs into male-specific brooding pouches located on their tail or abdomen (Wilson et al., 2001; Hoffman et al., 2006; Wilson, 2009; Winkler et al., 2012). Thus, fecundity in females is limited by abdomen volume, while in males brood size is a function of pouch size (Hoffman et al., 2006; Winkler et al., 2012). As expected, fecundity selection has been shown to target male brood pouches and female abdomens independently, thus causing clear region-specific sexual dimorphisms (Hoffman et al., 2006; Wilson, 2009; Winkler et al., 2012).

## V. THE FEMALE MULTIPLE-MATING COMPONENT OF FECUNDITY

Modelling the fitness dynamics of fecundity selection is a complex task given that fecundity is not simply a function of female traits that facilitate larger broods (e.g., body size), but a function of the pairing between males and females. Essentially, individuals do not have fecundity, but mating partners do (Pollak, 1978; Rice, 2004). Therefore, when modelling fecundity selection, fitness is assigned to mating pairs. In addition, the evolutionary dynamics of selection for fecundity rely on the tight dependence between selection for viability and for mating success, and hence, fecundity selection often reflects the interaction between natural and sexual selection (e.g., Badyaev & Ghalambor, 2001; Hoekstra et al., 2001).

Rice (2004) modelled the fitness dynamics of fecundity assuming mating scenarios represented by one female and one male only (see also Pollak, 1978). However, this assumption is a simplification of the dynamics of selection from variance in fecundity within populations, as it is broadly recognized that female multiple-mating (both remating and polyandry) influences fitness through increases in fecundity in a wide diversity of organisms, from invertebrates to vertebrates (Gwynne, 1984; Ridley, 1988; Madsen et al., 1992; Karlsson, 1998; Arnqvist & Nilsson, 2000; Eady et al., 2000; Evans & Magurran, 2000; Wiklund et al., 2001; Woolfenden et al., 2002; Kamimura, 2003; Vahed, 2003; Lewis et al., 2004; Torres-Vila et al., 2004; Bjork & Pitnick, 2006; Schwartz & Peterson, 2006; Engqvist, 2007; Gershman, 2007; LaDage et al., 2008; Lorch et al., 2008; McNamara et al., 2008b; Slatyer et al., 2012). This view is reinforced by the overwhelming evidence that female multiple mating is common in a wide diversity of species (Birkhead & Møller, 1998; Simmons, 2001; Cornell & Tregenza, 2007). Several studies suggest that this relationship is driven by a number of male traits that increase fecundity through increases in oogenesis and ovulation rates, such as nuptial gifts, courtship feeding, and nonspermatic beneficial components present in ejaculates (Simmons, 2001; Fedorka & Mousseau, 2002; Gillott, 2003; Poiani, 2006; Alonzo & Pizzari, 2010). Interestingly, these fecundity-stimulator traits are also known to play important roles in differential success in competition over mates. Therefore, the above observations suggest an adaptive basis for male-driven fecundity, and the complex evolutionary dynamics behind variance in fecundity as the result of interactions between sexual and fecundity selection. However, higher fecundity in polyandrous females has also been suggested to simply result from access to more sperm rather than to any fecundity-enhancing effects of male traits (Evans & Magurran, 2000).

The adaptive basis of the relationship between fecundity and multiple-mating has recently been reinforced by mathematical theory based on models assuming male and female prezygotic investment in offspring. Alonzo & Pizzari (2010) found that a strong effect of male fecundity-stimulator traits on female fecundity can shift the coevolutionary dynamics of female remating and male ejaculate expenditure from conflict to cooperation when mutual fitness is enhanced by increased fecundity. This model predicts that strong fecundity stimulation promotes cooperation between and within the sexes because a male fertilizes more eggs when mating with a promiscuous female. Therefore, when the costs of remating are compensated by the fecundity-mediated fitness benefits of promiscuity, selection for remating will prevail. However, in many other species, polyandry has negligible or no influence on fecundity enhancement. In these cases, conflict can be expected because the mating role of each male (i.e., first or second male precedence) will result in one of them taking advantage of the ejaculates of the other (e.g., Parker, 1998; Hodgson & Hosken, 2006), and because remating will compromise female’s fitness (Alonzo & Pizzari, 2010). Finally, when the intensity of selection on female remating avoidance opposes selection for remating because fecundity is not enhanced or because it compromises survival, males can cooperate among them while interacting in conflict with females (Alonzo & Pizzari, 2010). Consequently, this theory suggests that the interaction of fecundity with other components of fitness (e.g., survival) can result in cooperation over mating, in contrast to traditional expectations that sexual conflict necessarily arises from female remating.

## VI. FECUNDITY SELECTION ON MALES

Given that fecundity is predominantly limited by female capacity to produce ova and to provide embryos with the space they need for development, fecundity selection has historically been deemed as a female-specific force (Darwin, 1874; Williams, 1966; Roff, 2002; Fairbairn et al., 2007). However, despite this traditional view, a number of studies suggest that fecundity selection also operates on males when their phenotypes exert a direct effect on brood size (Savalli & Fox, 1998; Wilson, 2009; Winkler et al., 2012). Fecundity selection on males occurs primarily in species in which some extent of sex-reversed roles has evolved (i.e., any form of ‘male pregnancy’). In these organisms, eggs are transferred from the females into male structures specialized in retention of the embryos until they complete development (Cei, 1962; Kraus, 1989; Kolm, 2002; Winkler et al., 2012; Vitt & Caldwell, 2014). Cases are in fact abundant. For example, numerous amphibian species have evolved male ‘pregnancy traits’, such as back skin forming brood chambers that serve as nests (Greven & Richter, 2009; Fernandes et al., 2011; Vitt & Caldwell, 2014), inguinal pouches (Ehmann & Swan, 1985), or legs adapted to carry developing eggs (Marquez, 1993). Also, a quite unique case of male pregnancy is ‘neomely’. This strategy is exclusive to the Chilean frogs of the genus *Rhinoderma*, in which males have evolved vocal sacs where eggs are retained until they complete development, while the embryos are also provided with food (Cei, 1962; Goicoechea et al., 1986; Vitt & Caldwell, 2014). Similarly, a number of fish species show male body regions adapted for pregnancy. For example, paternal mouthbrooding provides embryos with ideal environments to complete development. In the species *Pterapogon kauderni*, Kolm (2002) reported that clutch size is a function of male body size. Thus, paternal phenotype plays a central role in fecundity in this fish. Also, in the *Syngnathus* pipefish, males have evolved specialized body pouches (on either the tail or abdomen) into which eggs are transferred by females to be fertilised and retained for development (Breder & Rosen, 1966; Wilson, 2009; Winkler et al., 2012). Interestingly, it has been shown in these fish that fecundity selection operates differentially on egg-brooding structures between the sexes, that the male skeletal structures that sustain brooding pouches have significant additive genetic variation, and that they are heritable and modular in the way they vary relative to other body regions (Hoffman et al., 2006). Similar cases exist in insects. For example, availability of male backspace in belostomatids (waterbugs), where egg-brooding takes place following transference from females, has been shown to directly limit female reproductive outcome (Kraus, 1989; Kruse, 1990).

A potential source of debate is that cases of male pregnancy can be interpreted as paternal care phenotypes driven by natural selection (e.g., Clutton-Brock, 1991) rather than by fecundity selection on males. However, it is important to distinguish between the origin of adaptations and their maintenance and variation once they have acquired a determinant role in reproductive success. Both phases of the evolution of adaptive traits can perfectly be targeted by different selective pressures. Thus, male pregnancy may originate via natural selection for paternal care, and subsequently be importantly shaped by fecundity selection once this trait has acquired an important role in fitness via differential fecundity. Indeed, this is likely to be the case for many traits subject to fecundity selection in general. For example, the evolution of reptilian viviparity is thought to be driven by natural selection arising from low temperatures on developing eggs (Hodges, 2004; Shine, 2005; Pincheira-Donoso et al., 2013). However, once viviparity has evolved in cold climates (where breeding seasons are considerably shorter), fecundity selection is predicted to counterbalance reduced opportunities for reproduction through increased fecundity (Cox et al., 2003; Pincheira-Donoso & Tregenza, 2011). Consequently, the roles of natural and fecundity selection on fecundity traits need not be mutually exclusive.

## VII. CONCLUSIONS

(1) Although fecundity selection has traditionally been treated as a force that influences fitness via increases in fecundity caused by larger body size in females, numerous studies have shown that this form of selection is not necessarily female-specific (e.g., it can target males in species with sex-reversed roles), and that it can also benefit fitness through reductions of brood size, or by targeting specific traits (e.g., abdomen) that may not result in larger overall body size. These expansions in the way fecundity selection operates in nature demand a redefinition of fecundity selection as a general mechanism that increases reproductive success through any effect on fecundity, caused by any trait, in either sex.
(2) Given that reproduction is energetically costly, the interaction between organisms and ecological pressures (e.g., food availability, predation) plays a central role in spatial and temporal variations in fecundity. As a result of both theoretical and empirical studies, it has become evident that predictable macroecological patterns of fecundity variation exist in nature (e.g., the widely known phenomenon we term Moreau-Lack’s rule). These patterns, which primarily express along latitudinal and elevational gradients, are explained by a number of ecological factors that vary with geography, such as seasonality, predation intensity, and opportunities for reproduction determined by length of breeding season. Although the macroecology of fecundity has consistently been shown, the overwhelming majority of research (especially studies testing for specific mechanisms driving the pattern) has focused on birds. Therefore, it is imperative to empirically assess the generality of these phenomena in a wider diversity of organisms. Both small- and large-scale studies are required in animals other than birds to quantify the contribution of the many potential factors suggested to affect fecundity along geographic gradients.
(3) Fecundity selection has, since Darwin (1874), been considered as one of the primary explanations for the evolution of female-biased SSD. However, given that fecundity selection often targets body regions that directly affect fecundity, and given that these traits may not necessarily affect overall female body size, we argue that the link between female-biased SSD and fecundity selection is not the generality. Indeed, it has been shown that strong fecundity selection can operate on females of species in which strong male-biased SSD exists. Finally, female-biased SSD can also result from male-specific selection for smaller body size. Therefore, we conclude that the action and strength of fecundity selection should not be estimated based on the extent of female-biased SSD, and therefore, that the absence of this form of SSD should not rule out the action and strength of fecundity selection.

## VIII. ACKNOWLEDGEMENTS

The authors thank Carl Soulsbury for a thorough critical review of the manuscript, which contributed to improve its angles and clarity. For important and insightful discussions on the topic, DPD thanks Tom Tregenza, Shai Meiri, Dave Hodgson and Dave Hosken. DPD thanks the University of Lincoln for a Research Investment Fund (RIF) Grant that supported this paper. JH was funded by NERC and a Royal Society Fellowship.

